# Electron microscopy reveals unique spore-like nano forms of *Bacillus cereus*

**DOI:** 10.1101/228833

**Authors:** Sumanta Ghosh, Biprashekhar Chakraborty, Shreya Ghosh, Sandip Dey, Chiranjit Biswas, Rukhsana Chowdhury, Krishnananda Chattopadhyay, Jayati Sengupta

**Author notes:** Contributed equally (Sumanta Ghosh, Biprashekhar Chakraborty). Corresponding Authors: Krishnananda Chattopadhyay & Jayati Sengupta. **Current Address:** NPDF Fellow, Department of Biophysics, Molecular Biology and Bioinformatics, Rajabazar Science College, University of Calcutta, India.

## Abstract

Endospore formation under environmental stress conditions is a well-established phenomenon for members of bacterial phylum Firmicutes, among which the most well studied ones belong to genus *Bacillus* and *Clostridium*. So far, known sizes of the spores are all larger than 500 nm. Nano-forms of bacteria have been reported but the notion still remains controversial.

In this study, we provide visual evidences of living nano-entities (named here as ‘nano-spores’) formed by a bacterial species *Bacillus cereus* under prolonged stress, which are capable of escaping though standard sterile filtration procedure. The existence of nano-forms of bacteria was initially identified in a yeast ribosome preparation. We further demonstrate the transformation of the ‘nano-spores’ into mature cells upon nutrient supply. Our study not only demonstrates the ability of bacteria to get transformed into yet-unknown form in order to survive under harsh environment, but also brings to light the existence of the smallest possible form of life.

## Introduction

Large environmental fluctuations induce transformation of a bacterial cell into its ‘starvation form^1^, which eventually gets converted into the ‘endospore’^2,3^. Bacterial endospores are considered the most resistant living form exhibiting incredible resistance to environmental extremities^4,7^. Under favorable growth condition, endospores germinate to vegetative forms eventually converting to mature bacterial cells^8^.

Although rod-shaped bacteria were initially identified as spore-forming species^9^, with emerging evidences it appears that a wide range of bacteria are capable of sporulation ^10–12^ Interestingly, it has been reported recently that in human microbiota, the spore-forming bacteria were significantly more diverse than the non-spore-forming bacteria^13^. While sporogenesis typically culminates in the production of a single endospore, there is evidence that some bacteria have evolved the means of producing more than one endospore per cell^14^

Based on recent structural studies of the endospore architecture of gram negative as well as gram positive bacteria^15,16^, it has been hypothesized that a spore is the common ancestor of all bacteria^17^. Emerging lines of evidence suggests that a wide range of bacteria are capable of sporulation^13,14^. The reported sizes of the endospores differ considerably^4,18^. Typically, mature spores are 0.8–1.2 μm in length and have either a dense spherical or ellipsoidal shape. The existence of nano form of life has been noted^1,6,7,19^, which has initiated a long-standing debate^11,20,21^.

Here, we provide evidences that *Bacillus cereus* can exist as nano scale structures (sizes of ~20-100 nm) which we have termed here as ‘nano-spores’ under prolonged stress. Interestingly, the nano-structures were first identified (and characterized as a form of bacterial species closely related to *Bacillus cereus*) in a yeast ribosome preparation. In marked contrast to previous reports on ‘nanobes’^7,19^, the ‘nano-spores’ that we have identified were obtained by subjecting *Bacillus cereus* cells to osmotic stress. The maturation of the nano scale structures back to mature bacterial cells was also tracked by exposing them to a nutrient rich environment.

## Results

### Identification of ultra-small spore-like structures camouflaged in purified ribosomes

Electron microscopic visualization of a preparation of yeast ribosomes revealing conspicuous small spherical structures (20-50nm) along with the ribosome particles attracted our attention (Fig. 1A). Intriguingly, when the yeast ribosomal preparation was incubated at 37°C with shaking for about 3 days, formation of larger particles was indicated by dynamic light scattering (DLS) analysis (Fig.S1A). Cell-like entities were also detected by atomic force microscopy (AFM) (Fig.S1C). Cryo-electron microscopy (cryo-EM) imaging revealed the presence of virtually transparent cell-like structures (0.6-1 μm) with double membranes (Fig.1B-D). These structures share striking morphological resemblance with the ultra-small bacterial cells recently identified^22^. Nano-scale spherical entities were also detected at the vicinity of the cell-like structures (Fig.1B-D insets). It must be emphasized that these cell-like particles were obtained only after incubation at 37°C, and they were not detected in the initial ribosomal preparation in spite of extensive search through several TEM grids.

**Figure 1:**
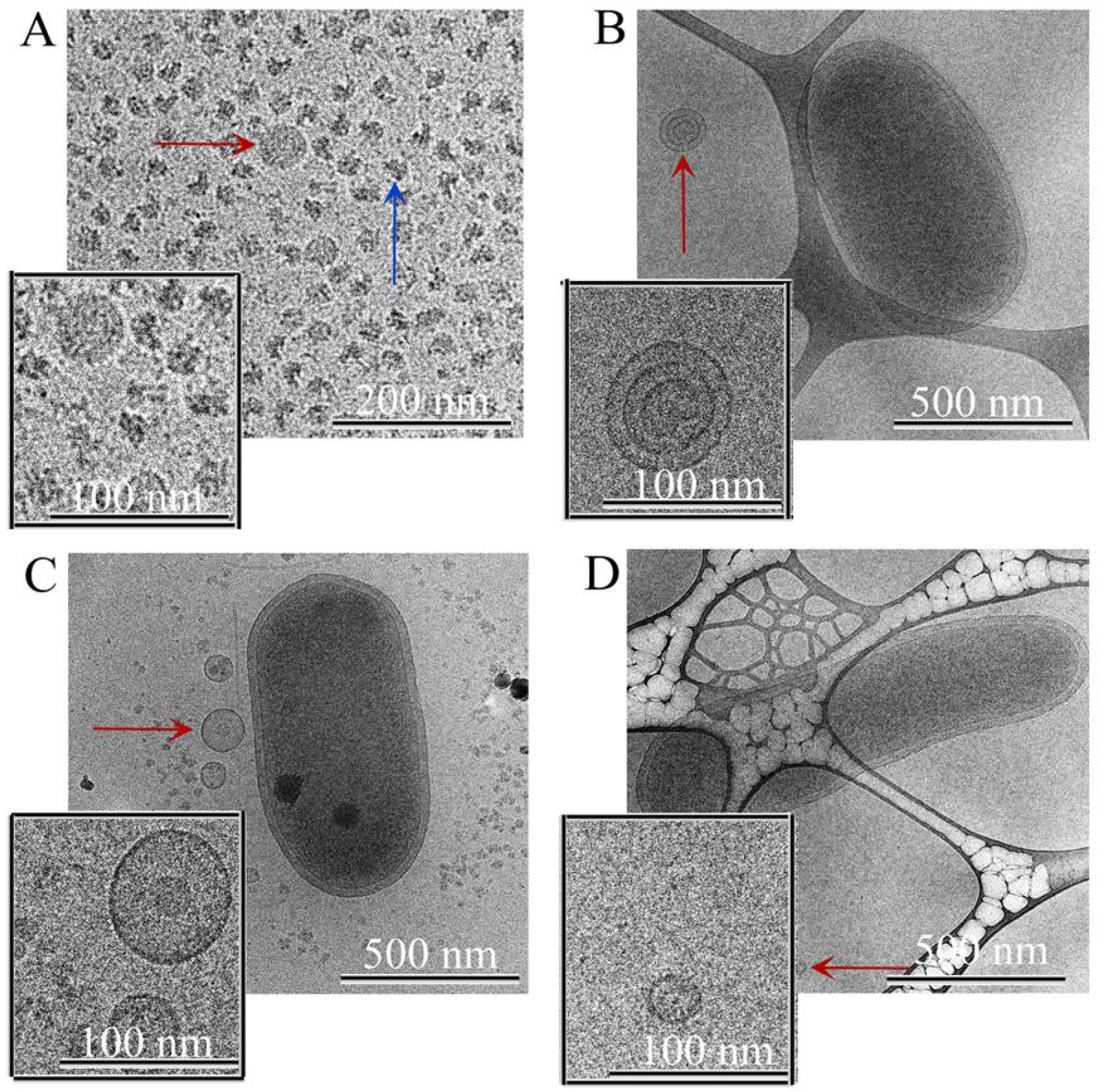
Cell-like structures observed in ribosome preparation. (A) A cryo-TEM image shows small spherical structures (marked with red arrow) along with ribosome particles (blue arrow) purified from yeast cells. (B-D) Large transparent cell-like structures (200 nm −1.5 μm) were detected in cryo-TEM when 80S ribosome preparation was incubated at 37°C for up to 72 hours. Nano-scale structures (red arrows) were observed in the vicinity of the large structures. Insets of A-D show close up views of nano-scale structures. Notably, ribosome particles were found to disappear with time as the time of incubation progressed.

DLS analysis of ribosome samples at different time points of incubation indicated gradual increase in size of the particles (Fig.S2E). Same samples, when visualized by cryo-EM, showed that the small spherical structures initially observed (Fig.1) grew in size with time to form cell-like structures (Fig.2A,C,E; Fig.S2A-S2D). It was further observed that addition of an intrinsically disordered protein during incubation expedited the enlargement of the cell-like structures (Fig. 2B,D,F; Fig. S1B, S2F). Interestingly, concomitant reduction in the number of ribosomal particles was observed over time (Fig. 2A,B).

**Figure 2:**
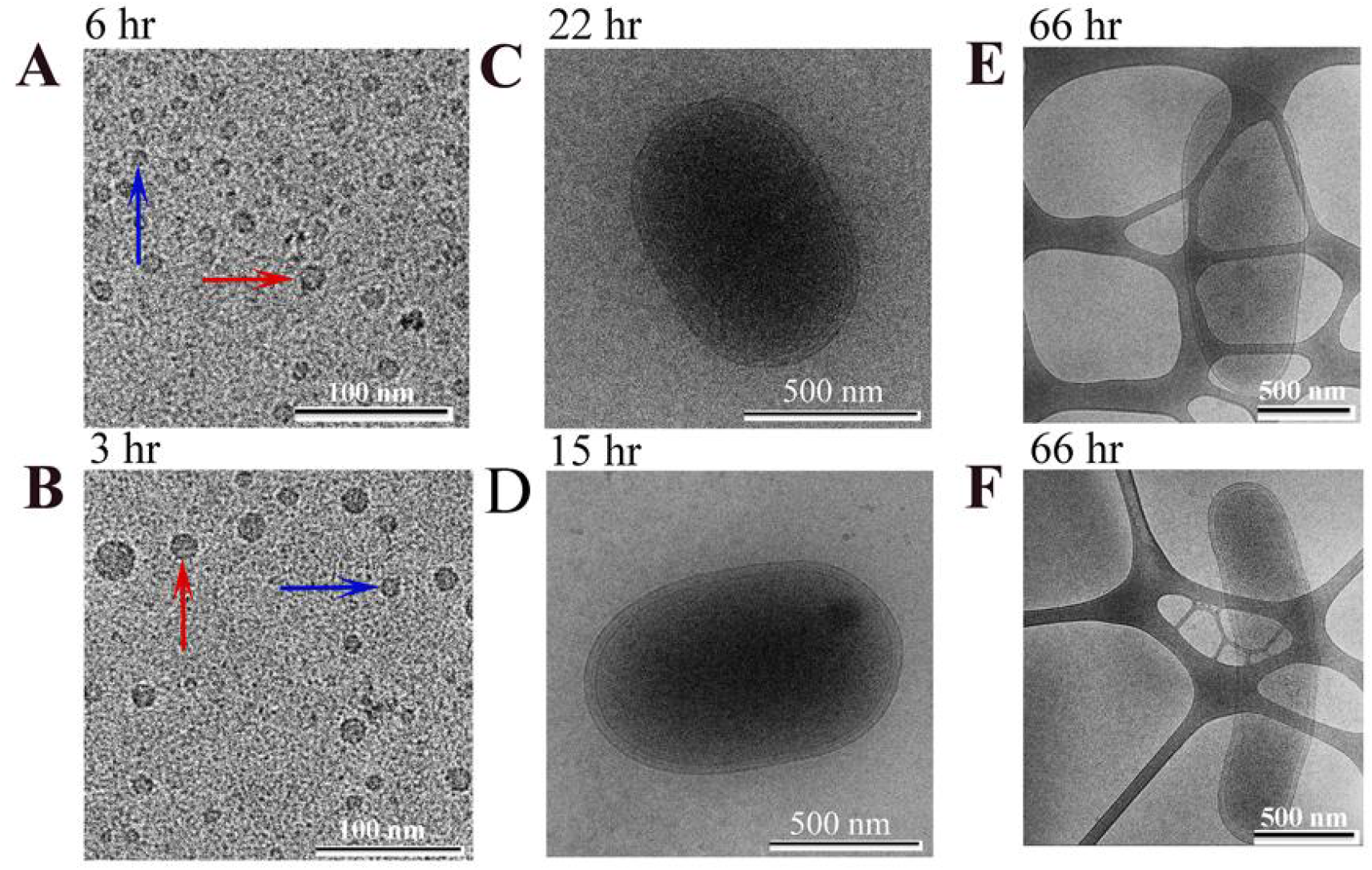
Tracking periodic changes of non-ribosomal particles in ribosome preparation. (A,C,E) Cryo-TEM images show the presence of nano-scale spherical structures (marked with red arrow) along with the ribosomes (blue arrow) after 6 hours of incubation of ribosomes at 37 °C. As the size of the small nano-scale structures (red arrows) increased with time, the number of ribosome particles simultaneously decreased. (B, D. F) The process described above was found to speed up in the presence of added intrinsically disordered protein, like alpha synuclein. (C) Large structures (500-800 nm) were observed after 22 hours of incubation of ribosomes, whereas, (D) particles of similar size were formed within 15 hours in the presence of externally added protein. At 66 hours, transparent cell-like structures formed in ribosome preparation (E). In presence of the protein (F) bigger size particles formed at similar time points.

We speculated that, during incubation at 37°C, ribosomal RNAs and proteins got disintegrated and acted as the nutrient source for the nano-scale structures to grow in size into larger entities (Fig. 2C,E). To confirm this assumption, we incubated the ribosome preparation with Alexa488 fluorophore-tagged α-synuclein protein (see Materials and Methods). While α-synuclein alone formed fibrillar structures after 48 hours (Fig. 3A), no fibrils could be detected when incubated in the presence of ribosome. In contrast, the fluorophore was found by confocal microscopy to be inside the cell-like structures (Fig. 3B) clearly suggesting that the protein molecules were engulfed by the nano-structures. Further, a RNA specific dye F22^23^ was applied to the ribosome preparation following 60 hours of incubation; subsequent confocal microscopy revealed the presence of the dye within the cell-like structures (Fig. 3C; Fig. S3A-C), which co-localizes with the protein (Fig. 3D,E), suggesting the presence of nucleotides within the cell.

**Figure 3:**
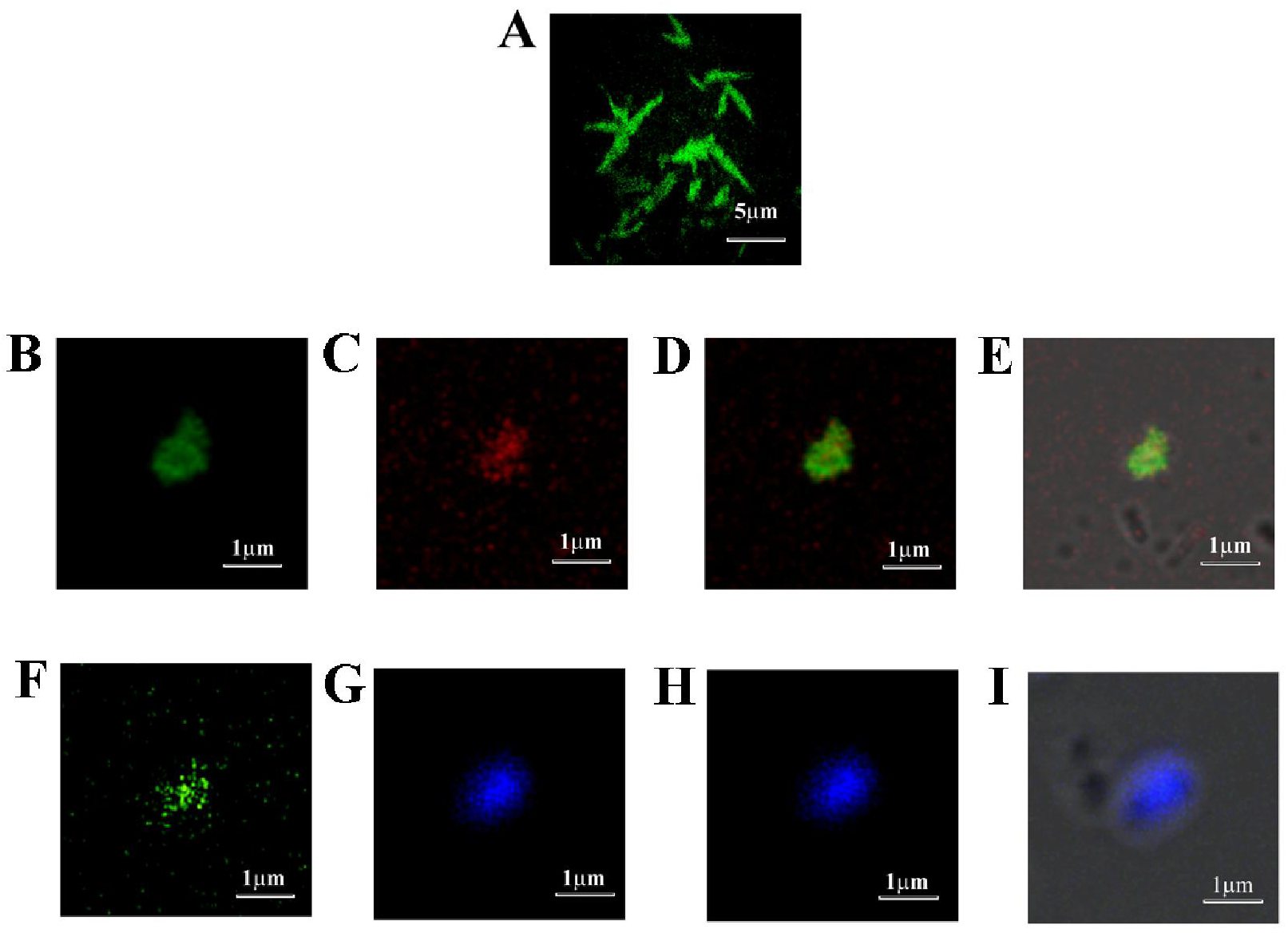
Localization of tagged-protein inside the cell-like living entity by confocal microscopy. (A) Confocal imaging showed fibrillar structures formed by the Alexa488 tagged α-synuclein (green fluorescence) when incubated at 37°C with shaking. (B) Fluorescence image following incubation of the protein along with yeast 80S ribosome preparation under similar conditions showed no fibrils; instead, some spherical structures of 05-1.5 um in size, detected by green fluorescence of Alexa488, co-localized inside. (C) Fluorescence imaging of the same region of interest (ROI) stained with RNA specific F-22 dye, also confirmed them as a living entities. (D,E) An overlay of green and red fluorescence and DIC images of ROI showed the green and red fluorescence to be localized within the cell-like structures. (F-I) Using Hoechst stain, fluorescence and DIC imaging showed colocalization of both protein and DNA inside such structures.

### Nano-scale spore-like structures identified in the ribosome preparation are living entities

The purified yeast 80S ribosome was spotted directly on Luria agar plates and incubated overnight at 37°C, along with a plate in which only the ribosome storage buffer was spotted as control. Unlike the control one (Fig.S4A), the plate with purified yeast 80S ribosome (Fig. S4B) showed distinct bacterial colonies confirming the nano-scale, spore-like structures, observed initially in the ribosomal preparation to be living entities. Considering the ability of the nano-scale spherical structures to convert into mature bacteria, we termed these nanobodies the ‘nano-spores’. However, in contrast to the conventional dense structures known for bacterial endospore, the ‘nano-spores’ detected here were almost transparent in nature (shown in Fig. 1).

We also performed confocal imaging of the ribosome sample, which was incubated with Alexa488 fluorophore-labelled α-synuclein protein (Fig. 3F) for 18 hrs using DNA-specific Hoechst dye. The images revealed the presence of the DNA-specific dye within the cell-like structures (Fig. 3G, Fig. S3D-F) co-localized with the protein (Fig. 3H,I), confirming the cell-like structures to be living entities. Further, DNA content analysis of the same sample at zero (0) hr and 18 hr of incubation, respectively, by flow cytometry revealed an increase in DNA content with time (Fig. S3G & S3H), suggesting that the ‘nano-spores’ are probably preparing themselves for germination into vegetative form.

Phylogenetic analysis of the universally conserved region of the16SrRNA gene, amplified from genomic DNA extracted from these bacterial cells, suggested that they belonged to the genus *Bacillus* (most related to *Bacillus cereus*; Fig. 4). However, the sizes of the cells were found to be much smaller (1-2μm) compared to the reported size of *Bacillus* and resembled the morphology of the forms of large spores or vegetative cells^24^. Conceivably, the *Bacillus* ‘nano-spores’ got co-purified with the yeast 80S ribosomes by virtue of their comparable sizes (~40-50 nm). Notably, association between *Bacillus* and yeast has been reported earlier^25^. Particularly, *B. cereus* has been shown to be capable of establishing parasitic or symbiotic association with fungi.^26,27^.

**Figure 4:**
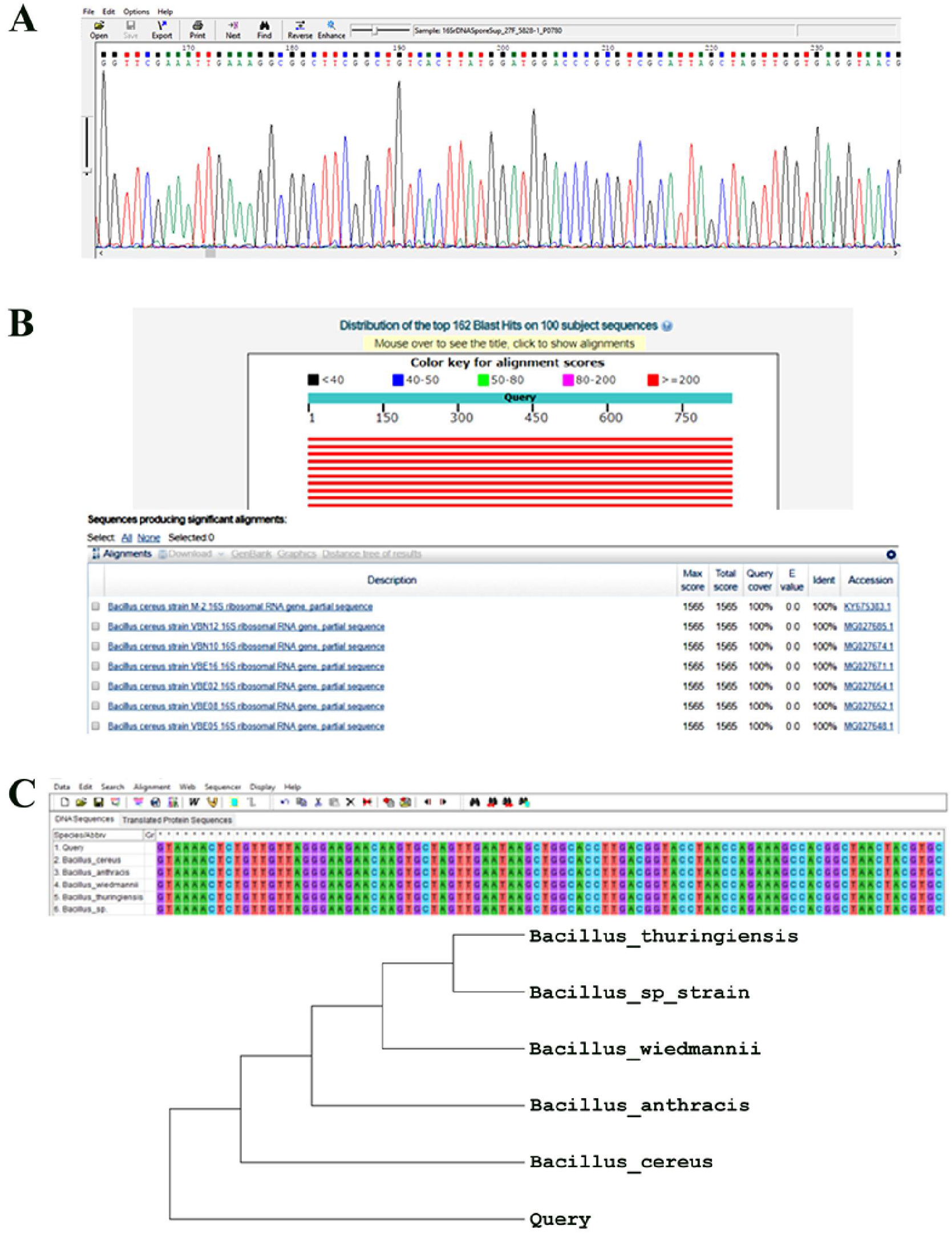
Phylogenetic identification of the bacteria by 16S rDNA sequencing. (A) Chromatogram of the “*query sequence*”, obtained from sequencing the 16s rRNA gene segment, amplified using universally conserved primers 27F and 1492R. (B) Nucleotide BLAST output of the *“query sequence”*, showing high sequence identity with different species of *Bacillus* genus. (C) Alignment of the *“query sequence”* with all the 16S rRNA gene sequences (from different *Bacillus* species), showing maximum sequence identity that was done using MEGA7. The phylogenetic tree was then constructed using “neighbor joining method” within MEGA7.

We examined the cultured bacterial cells under electron microscope. Remarkable network of pili-like structures, known to form in growing bacteria in order to exchange their genetic material, was observed (Fig. 5A,B) indicating its ability to rapidly grow under favourable growth conditions. In some views it appeared that the cells were embedded within a glutinous sack (Fig. S5A, B), which seemed to be a self-produced matrix resembling bio film architectures. Bio film formation has been reported earlier for various *Bacillus* species^4,12,18,28^.

**Figure 5:**
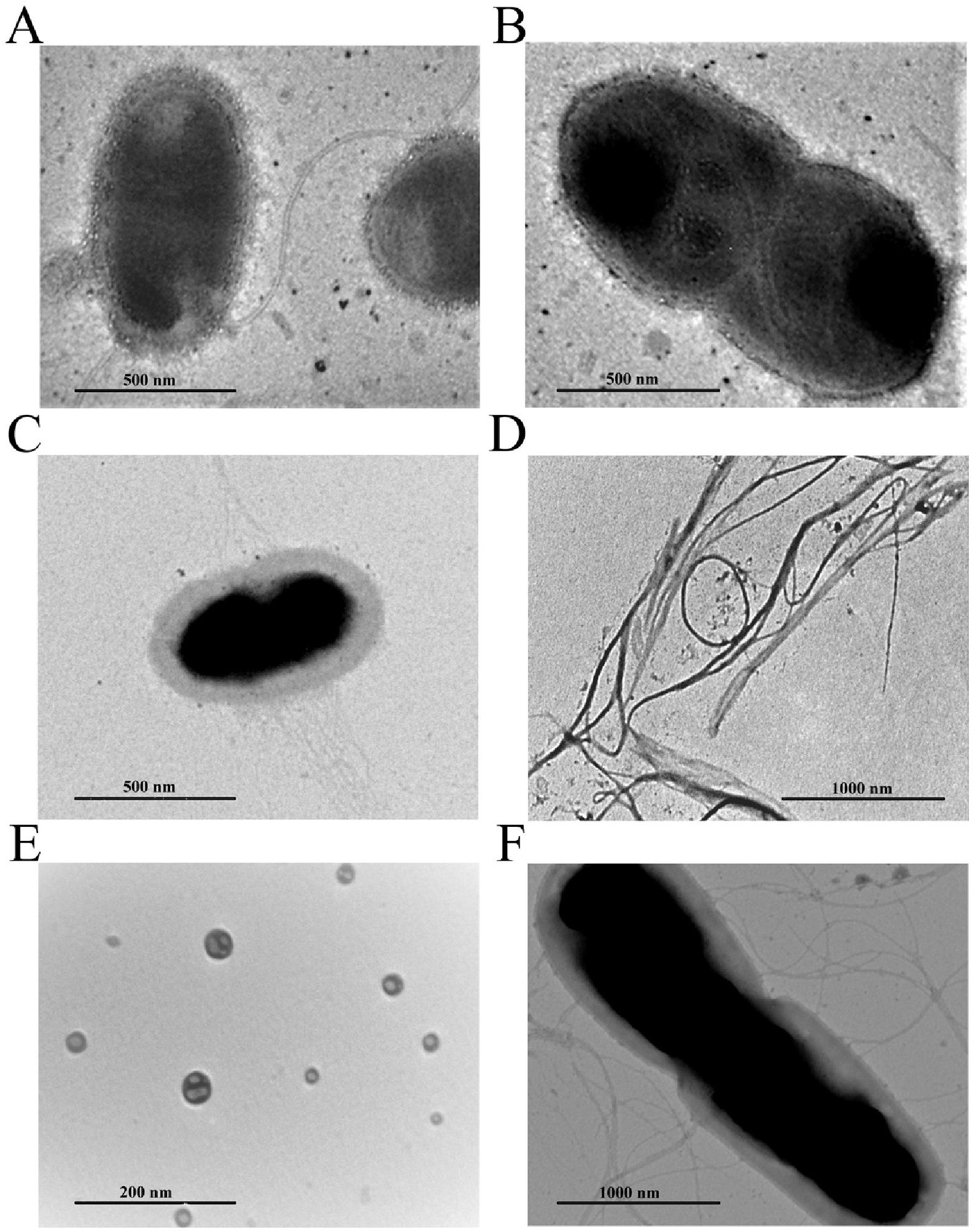
Confirmation of the presence of living entities. TEM images show (A, B) vegetative forms of bacterial cells when yeast ribosome preparation was cultured in Luria Broth (LB). Following overnight ethanol treatment, a precipitate was found at the bottom of the tube. The sample was subjected to centrifugation and (C) shrunken cells with fibrils coming off the structure were observed in the pellet, whereas (D) supernatant was full of fibrillar structures. (E) Nano-sized (20-50 nm) spherical structures (termed here as ‘nano-spores’) appeared in the supernatant upon prolonged salt-ethanol treatment and subsequent centrifugation. (F) TEM image shows growth of bacterial cells when the ‘nano-spores’ (formed under salt-ethanol stress) were allowed to grow in LB.

In line with an earlier observation^24^, the *Bacillus* strain in this study was found to be completely resistant to a number of antibiotics, like ampicillin and kanamycin, whereas a relatively high concentration of chloramphenicol (~70-140 μg/ml) and tetracycline partially inhibited bacterial growth (Fig. S5C, D).

### Efficient spore forming ability of the bacterial species

Bacterial cells are known to form highly resistant endospores during unfavourable environmental conditions. Hence, it may be assumed that mature bacterial cells (Fig. 5A, B), which were obtained from the ‘nano-spores’, can also be transformed to spores again, provided necessary external stress is applied. To verify this hypothesis, the cells were submerged in a solution of absolute ethanol and ammonium acetate (salt-ethanol treatment) and kept at −80°C (see online Methods). After overnight incubation, a precipitate was detected upon centrifugation, which was found to be shrunken cells when visualized under electron microscope (Fig. 5C). Unusual fibrillar structures were detected in the supernatant (Fig. 5D; Fig. S5E). Interestingly, a previous high-resolution AFM study reported that spore coat is composed of fibrillar structures^29^. Other studies also claimed that the amyloid fibrils form protective coat for the bacterial cells^30^. It is quite possible that under stress, amyloid-like fibrils (seen in supernatant) were released from the cell surface. It should be noted that *E. coli* cells lysed completely within overnight incubation when subjected to similar treatment (Fig. S5F).

The salt-ethanol treatment procedure consisted of multiple steps, which are described in detail in Fig. 6. Formation of the ‘nano-spores’ (~20-100 nm) was detected eventually in the fourth day supernatant when the sample was centrifuged (Fig. 5E). We also detected the presence of some particles, less than 200nm in size, containing unusual structural features (Fig. S6). The ‘nano-spores’ thus obtained (Fig. 5E) reverted back to vegetative forms when grown overnight in LB broth or in LB agar plate at 37°C (Fig. 5F; Fig. 6; Fig. S4C). 16S rRNA sequence phylogenetic analysis of these cells once again confirmed that this species belonged to the genus *Bacillus*. The whole cycle has been schematically represented in Fig. S7.

**Figure 6:**
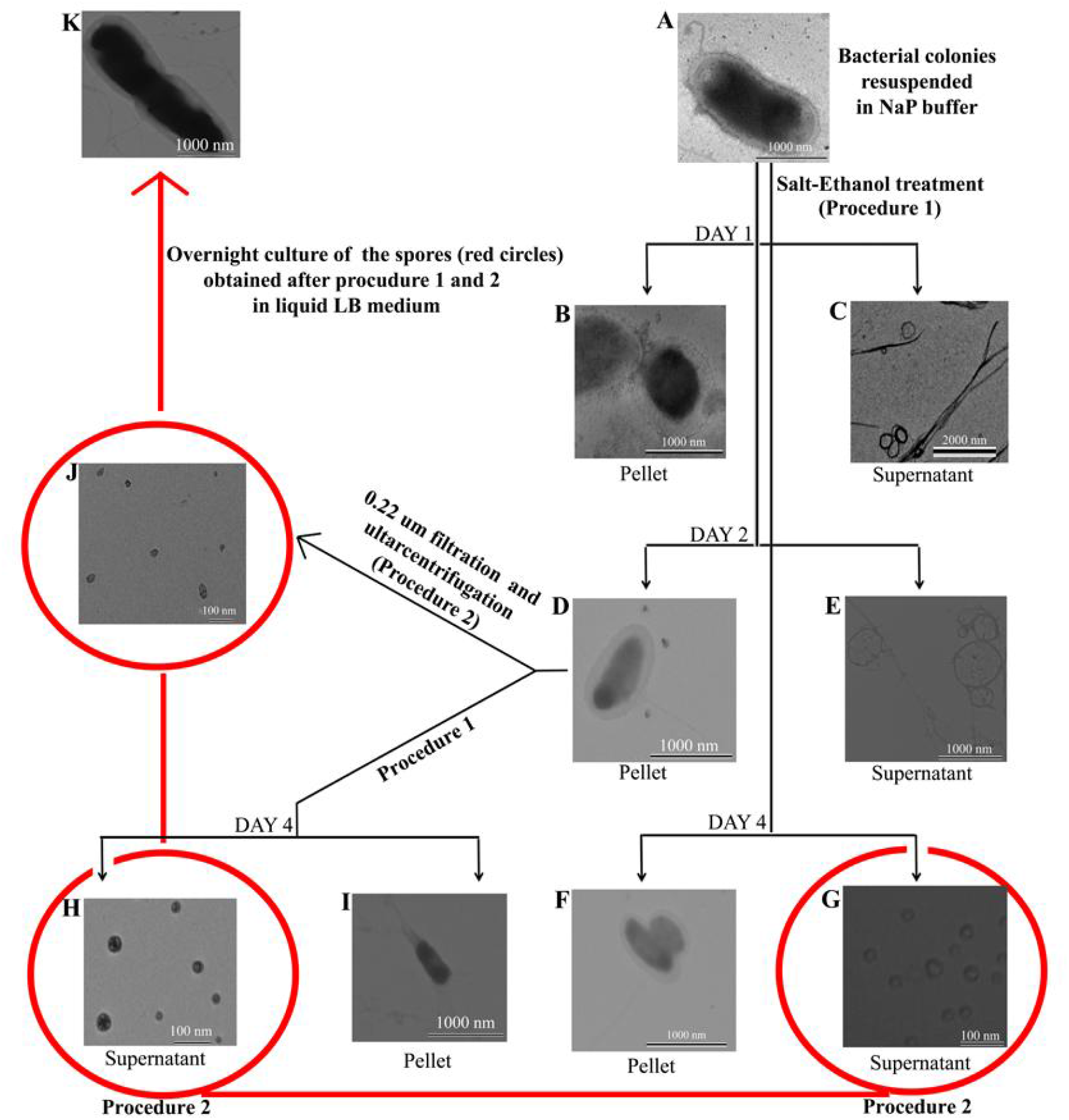
Detailed strategy to track nano-spore formation from bacterial cells under salt-ethanol stress. (A) TEM images of bacterial colony re-suspended in buffer. Centrifugation on first day following salt-ethanol treatment (marked as Procedure 1) showed (B) shrunken cells in pellet and (C) fibrillar structures in supernatant. After two days of treatment, (D) shrunken cells along with nano-scale spherical structures in the vicinity of the cells were observed in pellet, and (E) fibrillar structures were observed in supernatant as before. On the fourth day of treatment, upon centrifugation, supernatant and pellet fractions were collected. Supernatant fraction was again filtered through 0.22 micron syringe filter, followed by ultracentrifugation at the same speed used to precipitate yeast ribosome (marked as Procedure 2). (F) Few cells in pellet were seen, while, (G) ‘nano-spores’ (~20-80 nm) were detected in the supernatant fraction. The size of the cells (seen in pellet) gradually decreased from 1000-1200 nm to 800-1000 nm within 24 hours and further to 500-800 nm in 48 hours. Pellet obtained after two days of salt-ethanol treatment was separately subjected to further salt-ethanol treatment (Procedure 1), and on the fourth day supernatant and pellet fractions were collected following centrifugation. (H) ‘Nano-spores’ in the supernatant (following Procedure 2), and (I) shrunken cells in pellet were visible. The pellet obtained after two days of salt-ethanol treatment when subjected to Procedure 2, also led to the separation of (J) small size spores (30-60 nm) in solution. (K) Cultures of each of these ‘nano-spores’ (marked with red circles) on solid or liquid media, led to the formation of bacterial growth again.

Thus, we tracked the complete cycle of transformation of the bacterium, from ‘nano-spores’ (initially observed in our ribosomal preparation) to vegetative cells, and then back to ‘nano-spores’ by applying prolonged osmotic stress, which can again grow to mature cells (Fig. S7), confirming that the ‘nano-spore’ identified in this study are living entities.

### Bacillus cereus cells are capable of forming nano-spores under osmotic stress

Phylogenetic analysis indicated *Bacillus cereus* is the closest neighbour of the ‘nano-spore’- forming species identified in this study. Hence, we subjected *Bacillus cereus* M^1^_16_ strain (Fig. 7A) to similar salt-ethanol stress as mentioned earlier (Fig. 6). On the fourth day following salt-ethanol treatment (as mentioned previously in Fig. 6), both pellet (P) and supernatant (S) fraction were filtered through 0.22 μm filter (called fP and fS, respectively, henceforth) and subsequently viewed using cryo-EM. Nano-scale structures, very similar to those observed in purified yeast 80S ribosome were visualized in both supernatant (fS) and pellet (fP) (following removal of larger particles by filtration) fractions (Fig. 7B; 7E). More importantly, when the ‘fS’ fraction thus obtained was spotted on LB agar plate and incubated overnight at 37°C, bacterial colonies were observed (Fig. S4D), suggesting the nano-scale structures to be living entities.

**Figure 7:**
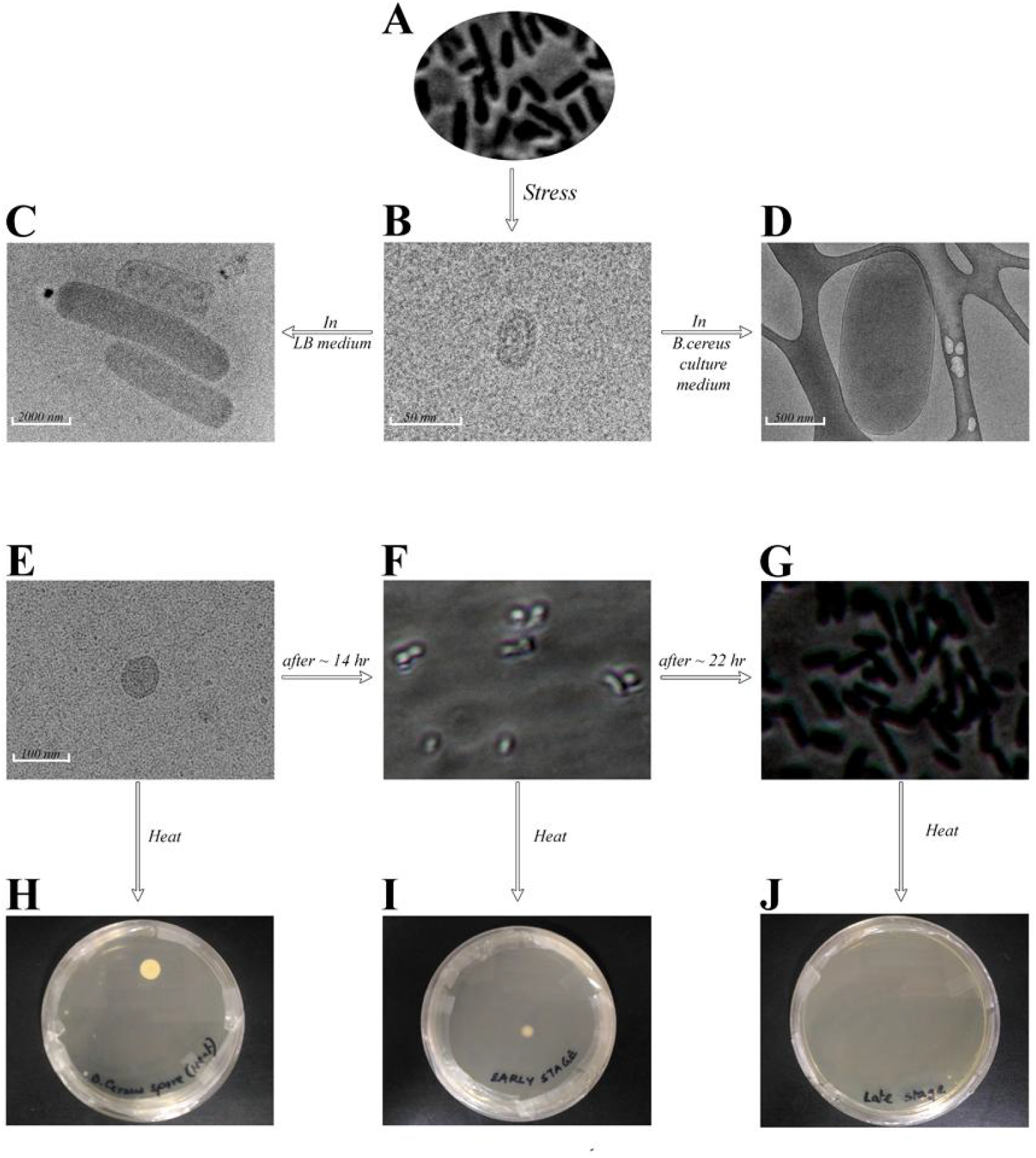
Characterisation of *B.cereus* nano-spores. (A) Mature *B.cereus* cells as seen under phase-contrast microscope. (B) Nano entities (termed as ‘nano-spores’) observed under Cryo-TEM after subjecting the *B.cereus* cells to osmotic stress. (C) Rod-shaped mature bacilli cell-like structures observed under Cryo-TEM after incubating the ‘nano-spores’ (observed in (B)) at 37°C with shaking in LB medium. (D) Transparent cell-like structures observed under Cryo-TEM after incubating the ‘nano-spores’ (observed in (B)) at 37°C with shaking in *B.cereus* minimal culture media. (E) *B.cereus* ‘nano-spores’ as observed under Cryo-TEM. (F) Ungerminated *B.cereus* spores observed under phase-contrast microscope after incubating the *B.cereus* ‘nano-spores’ at 37°C with shaking in LB medium for ~14hr. (G) Germinated *B.cereus* spores or cells observed under phase-contrast microscope after incubating the *B.cereus* ‘nano-spores’ at 37°C with shaking in LB medium for ~22hr. (H) Bacterial colony was observed after overnight incubation at 37°C from *B.cereus* ‘nano-spores’, as well as from ungerminated *B.cereus* spores (I) following heat treatment, but not from germinated *B.cereus* spores or cells (J).

It has been reported that supernatants of growing bacteria (containing muropeptides derived from the peptidoglycan fragments released by the bacteria) induce germination in dormant bacterial spores via a specific pathway^31^. All gram negative bacteria, as well as spore forming gram positive bacterial species, are known to release peptide fragments which can induce spore germination^31^. To see the effect of muropeptides released by *B. cereus* cells over the ‘nano-spores’ of the same species, we incubated both the ‘fP’ and ‘fS’ fractions in the supernatant obtained after harvesting an overnight *B. cereus* culture. Indeed, faster maturation of nano-structures was observed in the supernatant of *B. cereus* culture compared to its growth in sterile LB media, which is apparently due to the presence of muropeptides in the *B. cereus* culture media. Interestingly, cryo-EM images revealed a difference in appearance of the matured cell-like structures formed from the ‘fP’ and ‘fS’ fractions in sterile media (Fig. 7C) and *B. cereus* culture media (Fig.7D). While the cell-like structures in sterile media appeared dense, the structures formed in *B. cereus* culture media were more transparent, having striking similarity with those observed following incubation of yeast ribosome preparation at 37°C.

### Characterization of B. cereus nano-spores

Next, as we set out to track the maturation process of the *B. cereus* ‘nano-spores’ into mature cells, the ‘nano-spores’ were incubated in LB broth at 37°C with shaking (allows the enlargement of nano-forms as seen previously). Aliquots were taken at regular time intervals and visualized under phase-contrast microscope. Germination of a bacterial spore into a cell is known to be accompanied by a change in the refractility of the spore. As a result, spore loses its brightness, which makes the germinated cell appear dark under phase contrast microscope compared to an ungerminated spore^32,33^. Nano-structures are not visible in light microscope. Aliquots taken at an earlier stage of incubation (~14hr) revealed the presence of shining white (bright) particles (Fig 7F), while aliquots taken at a later stage (~22hr) revealed the presence of grey to black (dark) rod shaped particles (Fig 7G), similar to the B.cereus cells (Fig. 7A). Interestingly, an aliquot taken from an intermediate point (~18hr) revealed the presence of both bright and dark particles (Fig S4E & S4F), albeit segregated in separate microscopic fields. Our observations imply that maturation of *B.cereus* nano-structures occurs with an initial enlargement into the known forms of endospores (500-1000nm) followed by their germination into vegetative cells. It has been suggested earlier that a dormant but mature spore stores all the components necessary for germination^34^. Thus, the initial enlargement phase of the nano-structures (observed in our study) prior to germination is presumably required for getting themselves ready for germination.

The spores are known to be resistant to environmental extremities and some superdormant spores have also been reported having elevated level of resistance^35,36^. To test whether the nano-structures have similar kind of resistance, the nano-structures isolated following osmotic stress and an earlier stage (~14hr) aliquot from the previous experiment were kept in boiling water for half an hour and then streaked onto LB agar plate. Bacterial colony formation on a plate upon overnight incubation at 37 °C (Fig. 7H; 7I & 7J) is the evidence of a highly resistant nature of the nano-structures.

## Discussion

Existence of nano bacteria was claimed in the late 90’s^7,19^. However, this claim has particularly become controversial when it was found that non-organic materials, like calcium and phosphate ions, can hijack proteins from cell culture media to grow like nanobacteria. It was argued^21^ that the so called ‘nanobes’ are not ‘living units’, and could simply be fragmented portions of larger cells. We, for the first time, not only identified nano-scale, spore-like structures possessing fascinating characteristics but also provided unambiguous evidence that the structures are ‘living units’. Further, we discovered an unprecedented characteristic of *Bacillus cereus*, namely, to form nano-scale structures, and it probably won’t be too ambitious to suggest that many spore forming bacteria likely have the ability to form similar nano-structures.

The nano-size living entities we first identified were co-purified with the ribosome. To confirm that, all ingredients required for ribosome purifications were thoroughly checked to rule out any contamination of mature bacterial cells. Although we filtered the ribosome preparation through 0.22 micron filter prior to apply on agar plate, one far-fetched possibility was that the fully grown bacterial cells were co-purified with the yeast ribosome and got transformed to spores during storage (along with the ribosome) at −80°C. To examine that, we kept cultured bacterial cells (grown from the purified ribosome preparation) at −80°C in the same way as the ribosomes are preserved. The cells remained the same in size and morphology even after two weeks, ruling out this possibility.

Our results indicated that the nano-structures possess some of the spore characteristics. However, they may be considered as yet-another ‘starvation form’ of bacteria. It may be argued that nano-structures of this kind would be incapable of packaging essential biological macromolecules inside its small volume. It has been a matter of dispute whether macromolecules are essential for spore germination^8,33,34,37–40^. The prevailing view was that the spores can go on to germination without the need of any macromolecular synthesis, which starts only after germination, as the germinated spore gets converted into a growing cell^8,34^. However, this idea was challenged by Sinai *et.al* arguing that protein synthesis is necessary for spore germination^33,37,39^. Although the debate is still ongoing^39^, it is now believed that storage of the bio-macromolecules is not essential for germination, and that these molecules can be generated rapidly just before germination^39^. Our observation of the enlargement of the ‘nano-spore’ size, as well as an increase in DNA content prior to germination supports this notion.

It is evident from our study that ‘nano-spores’, by virtue of their size similarity with ribosomes, can get purified together with the organelles. However, it is not clear whether those are engulfed inside the yeast cells or if they remain attached to the cell surface. Nevertheless, the ribosome preparation protocol eliminates cell membrane and other debris at the very first step, and, thus, we suspect that the ‘nano-spores’ likely hide inside the yeast cells. Apparently, other microbes outnumber them in normal environment and, hence, they prefer to stay in ‘nano-spore’ form unless favourable condition is available^8^. The small size of the spores, which would make a typical sterile filtration process ineffective, and their persistence even after salt-alcohol treatment, requires serious attention, e.g., in case of hospital-acquired antibiotic resistance and microbial survival in extraterrestrial space.

## Methods

### Ribosome Purification

A diploid wild type strain MATaα derived by combining two haploid strains, 8534-10A (MATa, leu2, ura3, his4) and 6460-8D (MATα, met3), of *S. cerevisiae* was grown overnight in Yeast-Peptone-Dextrose (Y.P.D.) complete medium at 30°C. After around 18 hours the cells were harvested at 6000g for 10 min at 4°C. The cell pellet was re-suspended in ribosome buffer (100mM KOAc, 20mM HEPES-KOH, pH 7.6, 10mM Mg (OAc)_2_, 1mg/ml heparin, 2mM DTT, 0.5mM PMSF) and lysed by passaging twice through French pressure cell (16000 psig at first and then 18000psig). Lysate was centrifuged at 17000g for 30 mins at 4°C. Supernatant was then subjected to ultra-centrifugation at 350000g for 1hr at 4°C. The pellet obtained after centrifugation was re-suspended in minimum volume of ribosome buffer and the volume was made up to18ml by adding high salt buffer (ribosome buffer plus 500 mM KCl) and kept on ice for 60 min with gentle stirring at regular intervals. This was followed by centrifugation at 16000g for 10 min at 4°C for 3 to 4 times until no pellet could be visible. The supernatant was layered over a sucrose cushion and centrifuged at 350000g for 1 hr at 4°C. Pure yeast 80S ribosome obtained as pellet was re-suspended in storage buffer (10mM Tris-HCl (pH 7.5), 12.5mM Mg(OAc)_2_, 80mM KCl, 5mM 2-Mercaptoethanol, 0.5mM PMSF) and stored at 80°C.

Considering ribosome purification protocol, it is quite unlikely for mature bacterial cells (more that 10 fold differences in their sizes) to be co-purified with ribosome. Instead, it is more likely that the bacterial spores might have co-purified with the yeast 80S ribosome probably by virtue of their comparable sizes (yeast ribosomes ~40-50 nm).

### Transmission Electron Microscopy

Electron microscopy images were obtained with a Tecnai^™^ G2 Spirit transmission electron microscope at an acceleration voltage of 120 kV. A drop of the sample solution (4 μl) was spotted on a carbon-coated copper grid. The grid was dried carefully, keeping it in a vacuum desiccator for 24 hr and was finally examined under the electron microscope to study the ultra-structural features of the sample (bacterial cells/spores) at different conditions. Cryo-electron Microscopy (Cryo-EM):

Cryo-grids were prepared with standard protocol^41^. Ribosome preparation following incubation at different time points of incubation was spotted on glow-discharged lacey carbon copper grid and plunge-frozen in liquid ethane using Vitrobot (FEI Company). Grids were transferred to a cryo-holder and visualized in Tecnai POLARA (FEI Company) equipped with 300 kV field emission gun (FEG) at low electron dose. Images were collected at 82600× magnification (~5 μm under focus). Images were captured on an Eagle 4k × 4k CCD camera (FEI Company) with a final pixel size of 1.86 Å.

### Atomic Force Microscopy (AFM)

AFM was performed using a Pico Plus 5500 AFM (Agilent Technologies, USA) with a Piezo scanner having a maximum range of 9 μm. After incubating for 44 hours, 10 ul of newly formed cells was deposited on a freshly cleaved muscovite ruby mica sheet (ASTM V1grade ruby mica from MICAFAB, Chennai). After 30 minutes the sample was dried with a vacuum drier. The cantilever resonance frequency was 150-300 kHz. The images (256 pixels × 256 pixels) were captured using a scan size of between 0.5 and 8 μm at a scan speed of 0.5 lines/s.

The length, height, and width of protein fibrils were measured manually using PicoView1.10 software (Agilent Technologies, USA).

### Dynamic Light Scattering (DLS)

The Malvern Zetasizer Nano ZS DLS system (Malvern Instruments Ltd., UK) was used to perform all DLS measurements. This system is equipped with a 633 nm He-Ne laser and an APD detector configured to collect backscattered light at 173°. The measurement sample was allowed to equilibrate for 120 s at 20°C prior to the measurements. Fifteen separate runs, each of 10s duration were averaged for each DLS measurement. DLS data were analyzed using Malvern Zetasizer 6.12 software. The reported mean particle hydrodynamic radii (R_H_) were calculated using the intensity based particle size distributions.

### Confocal Microscopy

For the assessment of protein consumption inside the bacillus cells, we fluorescently labelled both protein and ribosome with specific dyes. The protein α-synuclein was labelled with Alexa fluor 488 C_5_ maleimide (alexa488) using standard protocol^42^. After labelling we did vigorous dialysis followed by size exclusion chromatography to remove all the free dyes in solution. The labelling efficiency of α-synuclein (α-syn) was 20%. Mature fibrils are formed after agitation of 200nm labelled α-syn (Total concentration of α-syn is 200μM). Excitation and emission wavelength of alexa488 is 488nm and 520nm (blue fluorescence) respectively.

Existing RNA was labelled by F-22 RNA dye following published procedure^23^. Excitation and emission wave length of F-22 dye is 548nm and 620nm (red fluorescence) respectively. For DNA identification, cells were labelled with Hoescht 33342 (Thermo Fisher Scientific). The excitation and emission wavelength is 361 nm and 497 nm. After labeling, the cells were observed under a Leica DMI6000 B inverted microscope equipped with Plan Apo 100X/1.40 oil objective (Leica TCS SP8 confocal system) and Leica DMI6000 B inverted microscope equipped with Plan Apo 100X/1.40 oil objective (Leica TCS SP8 confocal system).

### DNA Count

For DNA content in solution with increasing hours of incubation we stained the cells at 0 and 18 hours with DNA specific Hoescht 33342 dye and quantified the DNA count using BD LSRFortessa FACS instrument. Fluorescence intensity of Hoescht dye was measured at indo-violet channel with excitation wavelength 355 and emission range 425-475nm. The data were analysed by BD FACS Diva software.

### Salt-ethanol treatment on the bacterial cells

Ribosome preparation was directly applied on LB agar and kept at 37°C. Upon overnight incubation slimy bacterial colonies were observed. The bacterial colonies were re-suspended in LB broth and incubated at 37°C with shaking. Log phase cells were harvested and resuspended in 20mM sodium phosphate (pH 7.5) and then half the volume of 7.5M ammonium acetate was added to it. This was followed by addition of double the total volume of absolute ethanol and kept at-80°C. Similar treatment was carried with *Bacillus cereus* cells.

Salt-ethanol treated cell suspension was subjected to centrifugation at 13000 rpm at different time points yielding pellet and supernatant fractions. The pellet obtained was further resuspended in sodium phosphate buffer (pH 7.5) for further investigations. Notably, when similar treatment was applied overnight to *E. coli* cells (control bacterial species) no pellet was detected upon centrifugation and only cell debris was observed in the supernatant. The ‘nano-spores’ were observed at different steps of treatment in both supernatant and pellet fraction. Such fractions containing ‘nano-spores’ were filtered through 0.22 um filter followed by ultracentrifugation at 350000 g (the speed at which yeast 80S ribosome is precipitated) to precipitate the nano-spores.

### Determination of spore character of the nano-spores

Spore suspensions fS and fP (200 μl) were added to 1 ml growth media and incubated at 37°C with shaking. At specific time intervals aliquots were taken for heat-treatment and microscopy.

### Phase-contrast microscopy

Samples were air dried, heat or mathol fixed on glass slides and visualized by positive phase contrast microscopy under 100X oil-immersion objective lens, phase-3 position in an upright bright field-phase contrast microscope (DM-2500, Leica Microsystems, Germany). The viewer was unaware of the expected outcome of the respective stages of the samples to avoid any biasness. An additional 2.5 digital zoom factor was incorporated for higher magnification.

## Acknowledgements

This work was supported by CSIR-Indian Institute of Chemical Biology, Kolkata, India, CSIR Network project ‘UNSEEN’ and SERB, DST (India) sponsored projects. SG and SD acknowledge UGC, India, for senior research fellowship. We would like to thank Prof. Syamal Roy for helpful discussions. We also thank Dr. Anirban Kundu for helping in phylogenetic analysis, Mr. Diptodeep Sarkar Mr. Binayak Pal, and Mr. Somnath Mazumder for technical help with microscopy, Mr. Tanmoy Dalui for FACS measurement, Prof. Young-Tae Chang, Department of Chemistry, National University of Singapore, for gifting us the RNA specific F22 dye and Dr. Animesh Naskar for gifting us the *Bacillus cereus* strain.

## Author contributions

SG(1) and BC performed experiments to monitor bacterial transformation to spore and spore to mature bacteria, 16S rRNA preparation for sequencing. SG(2) assisted SG(1) and BC to perform experiments with *B. cereus* cells. SD performed antibiotic experiments. RC analyzed the 16S rRNA gene sequencing data and shared expertise to critically check the results. CB performed TEM and cryo-TEM surveys, KC and JS conceived the project, supervised research and JS, BC and KC wrote the paper with inputs from SG(1).

