## Supporting Information for "Electron microscopy reveals unique spore-like nano forms of *Bacillus cereus*"

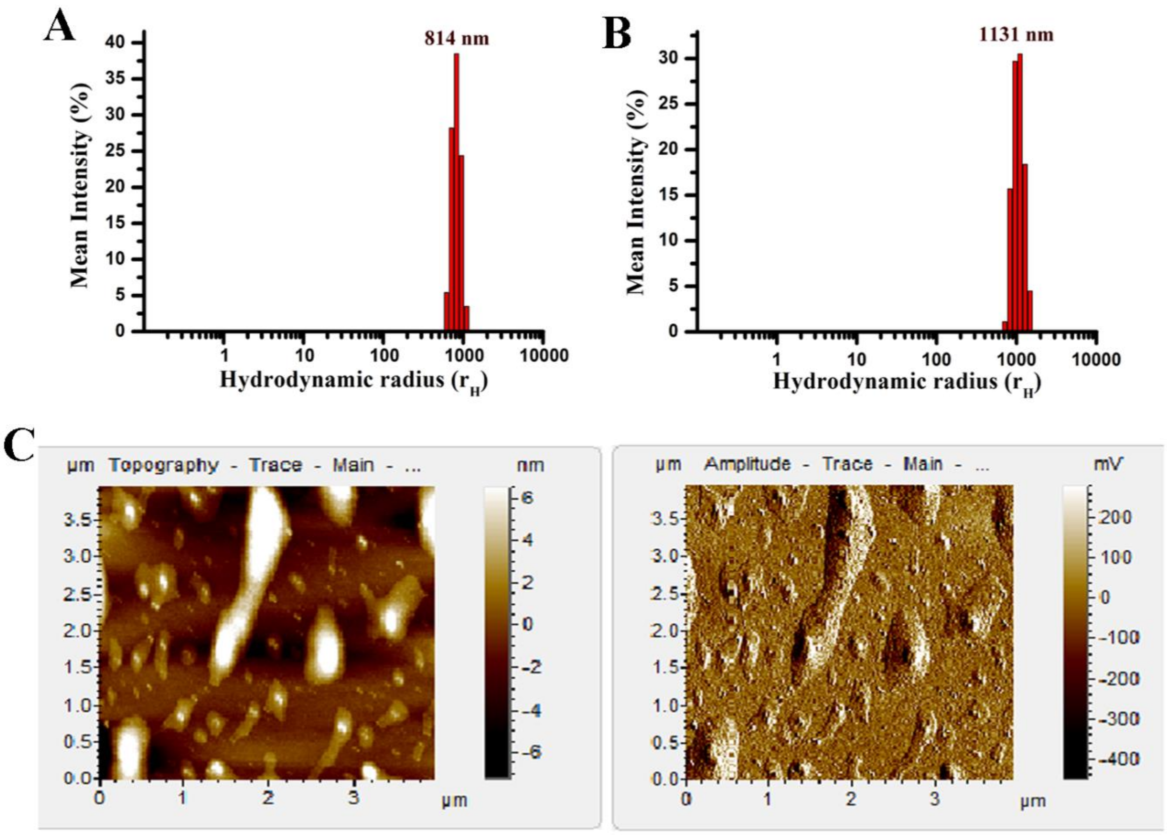


**Figure S1. Formation of large particles upon incubation of the ribosome preparation at 37°C with shaking.** (A) Dynamic Light Scattering (DLS) measurement also confirmed the presence of large particles in solution when ribosome preparation was incubated for long time. (B) DLS measurements also showed that particles formed in presence of disordered protein were larger in size at similar time point. (C) Atomic Force Microscopy (AFM) revealed the presence of large particles having cell-like features.

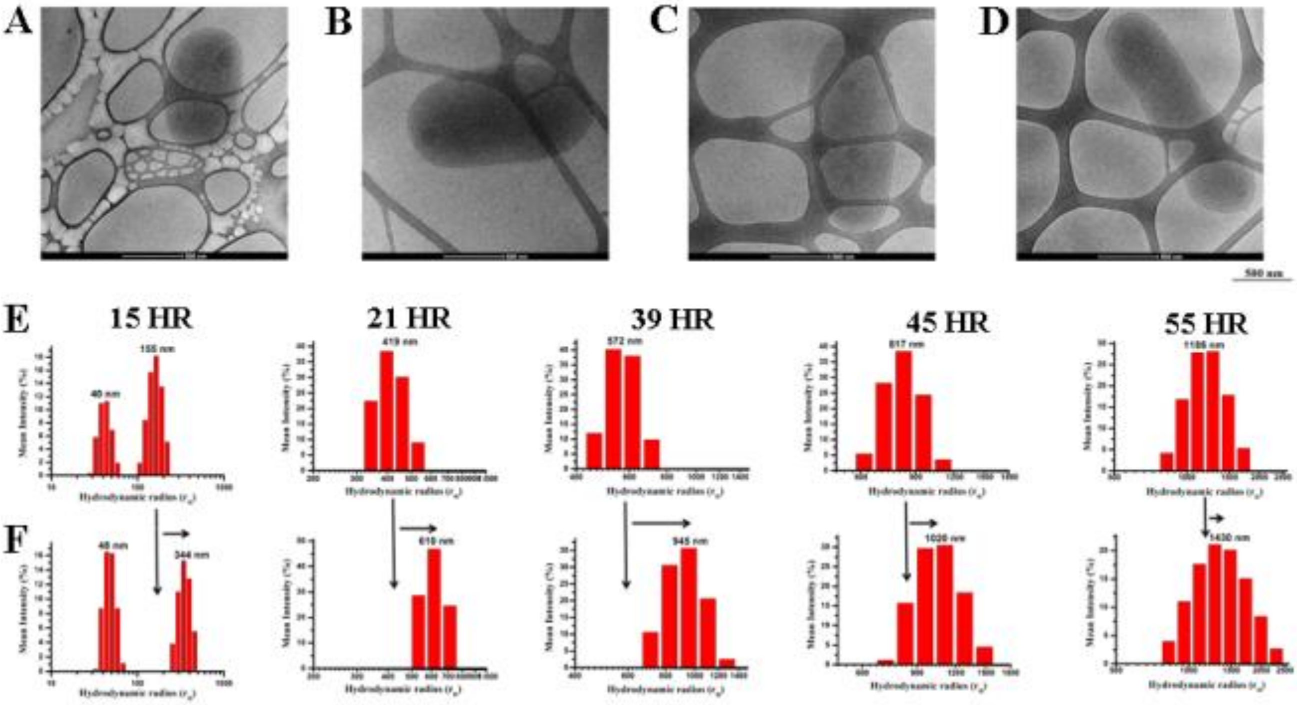


**Figure S2: Comparative analysis of enlargement of the particles with time.** (A-D) Cryo-EM images show that the size of the cell-like particles increases with time when the ribosome preparations were subjected to incubation at 37$^{o}$C for hours. DLS measurement detected formation of larger particles when only ribosomes were incubated (E), and when the protein (A-syn) was present during incubation (F) at different time points (15, 22, 39, 45 and 55 hours). At specific time point, there is a right shift in peak of hydrodynamic radius (shown by black arrow) with addition of protein suggesting that change in size was more rapid when protein was present.

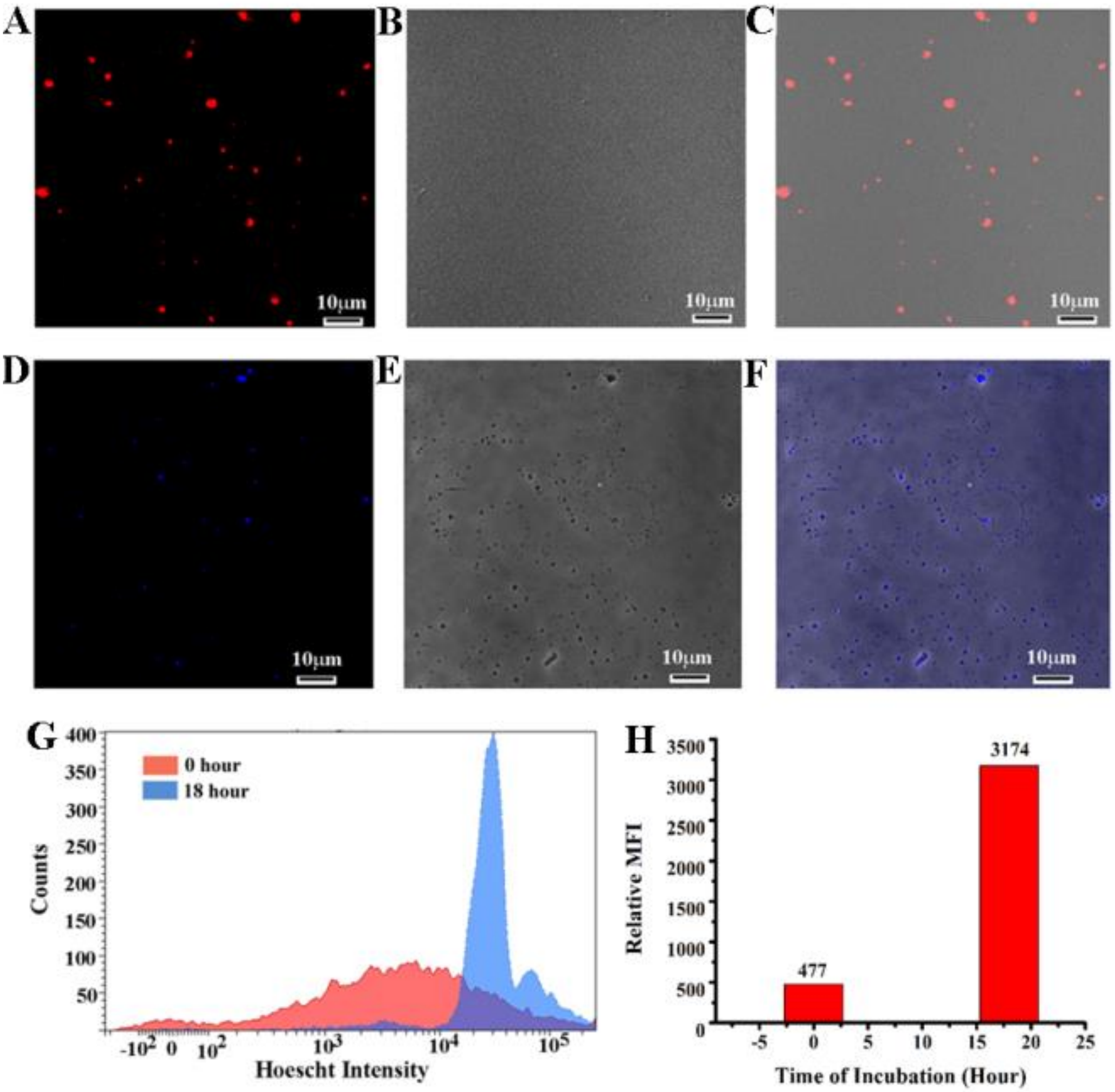


**Figure S3: Localization of tagged-DNA and RNA inside the cell-like structures by confocal microscopy.** (A) Using RNA specific dye F22, confocal microscopy image of the ribosome preparation after 39 hours incubation at 37$^{o}$C with shaking detected identical spherical structures by virtue of the red fluorescence of the dye. (B) DIC imaging of the ROI identified cell-like structures, and upon merging (A) and (B) the red fluorescence was found to be localized within the cell-like structures (C). (D) Fluorescence image of the ribosome preparation after 18 hours incubation with DNA specific dye Hoechst at 37$^{o}$C with shaking, detected spherical structures of 0.5-1.5 µm size by virtue of the blue fluorescence of the dye. (E) DIC imaging of the ROI identified cell-like structures, and upon merging (D) and (E) the blue fluorescence was found to be localized within the cell-like structures (F). (G) Distribution of Hoechst intensity by FACS after 0 hour and 18 hours of incubation of ribosome. (H) Relative mean fluorescence intensity (MFI) of Hoechst at different time (0 and 18 hours) has also been shown.

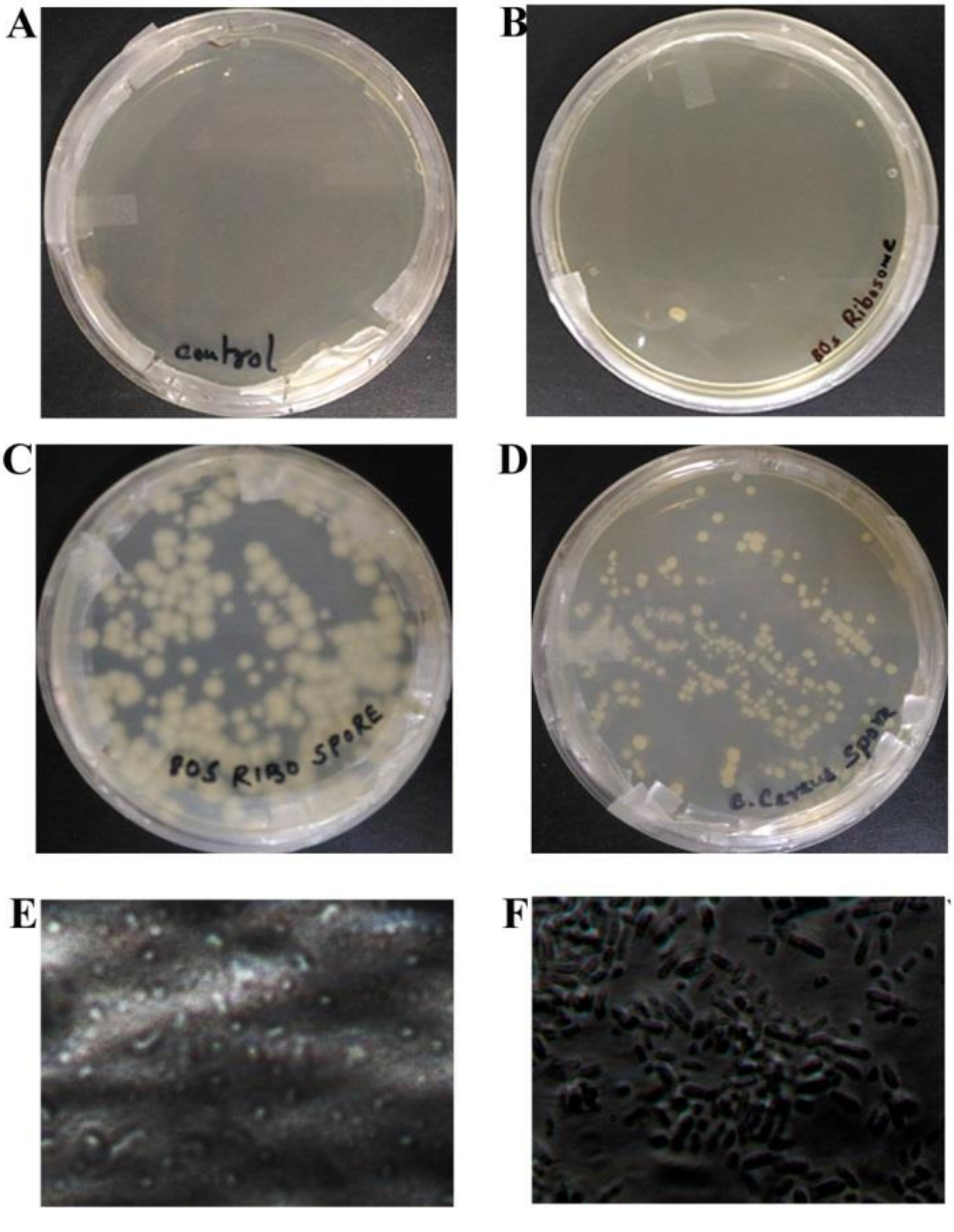


**Figure S4: Nano-spore Characterisation.** (A) Control LB agar plate, spotted with ribosome storage buffer and incubated at 37$^{o}$C overnight. (B) LB agar plate, spotted with purified yeast 80S ribosome preparation and incubated at 37$^{o}$C overnight. (C) LB agar plate, spotted with the nano-spores produced by reverting back the bacterial cells (which were obtained from maturation of the nano-entities in yeast 80S ribosome preparation) under salt-ethanol stress and incubated at 37$^{o}$C overnight. (D) LB agar plate, spotted with the nano-spores produced (under salt-ethanol stress) from the *B.cereus* cells and incubated at 37$^{o}$C overnight. (E) Shining white ungerminated *B.cereus* spores as well as dark germinated *B.cereus* spores or cells (F) as observed in different fields under phase-contrast microscope after incubating the *B.cereus* nano-spores at 37$^{o}$C with shaking in LB medium for ~18hr.

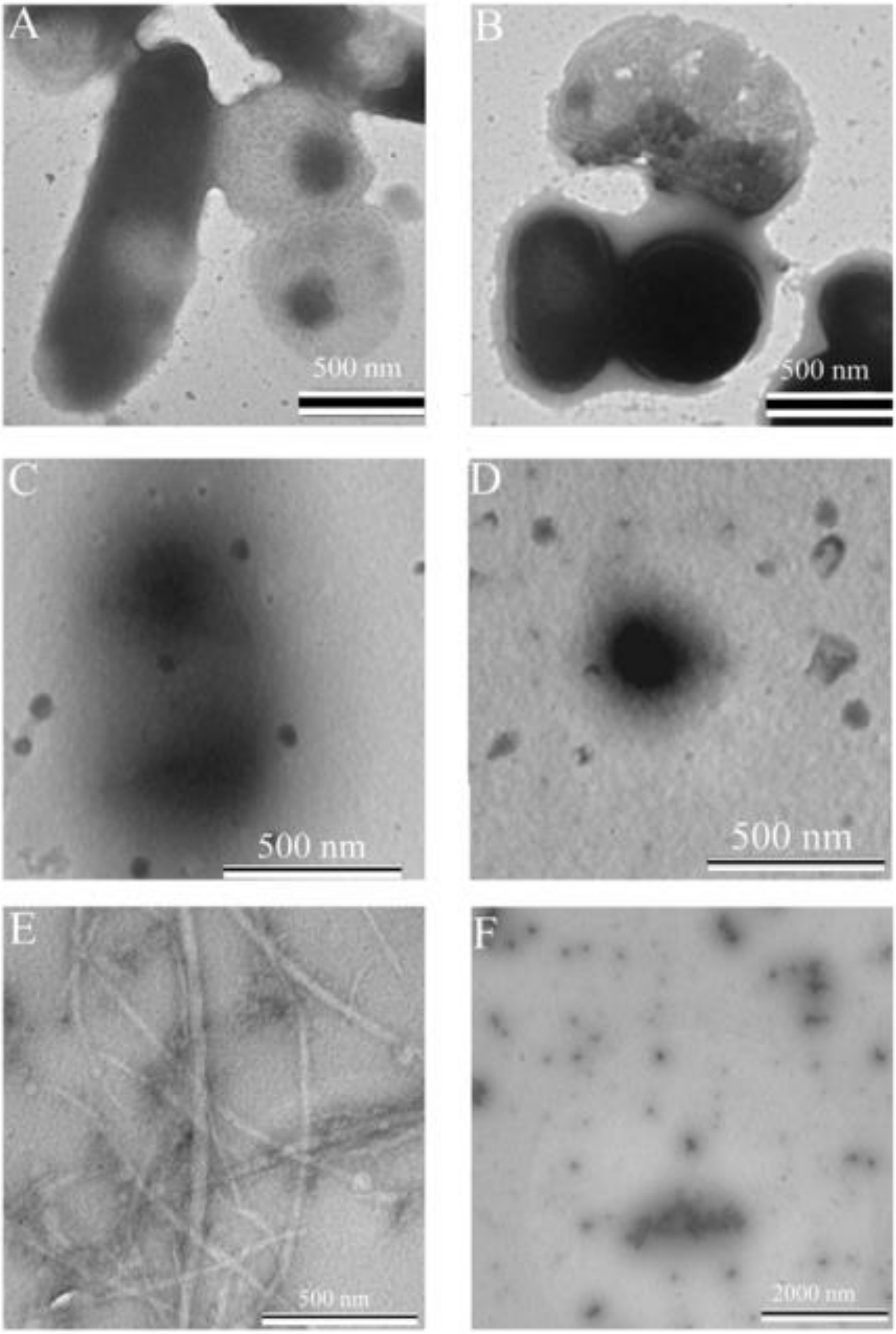


**Figure S5: Bacterial cells and effects of different treatments.** (A-B) Additional cellular features of cultured bacterial cells re-suspended in phosphate buffer and visualized under electron microscope (120 kV TEM). (C) Higher doses of chloramphenicol and (D) tetracycline arrested the growth of the bacteria and destroyed the cells, as seen in TEM images. (E) Features of fibrillar structures observed in the supernatant following vigorous treatment with salt-ethanol. (F) *E. coli* cells grown in LB medium following treatment with salt-ethanol - no pellet was detected and supernatant contained only debris.

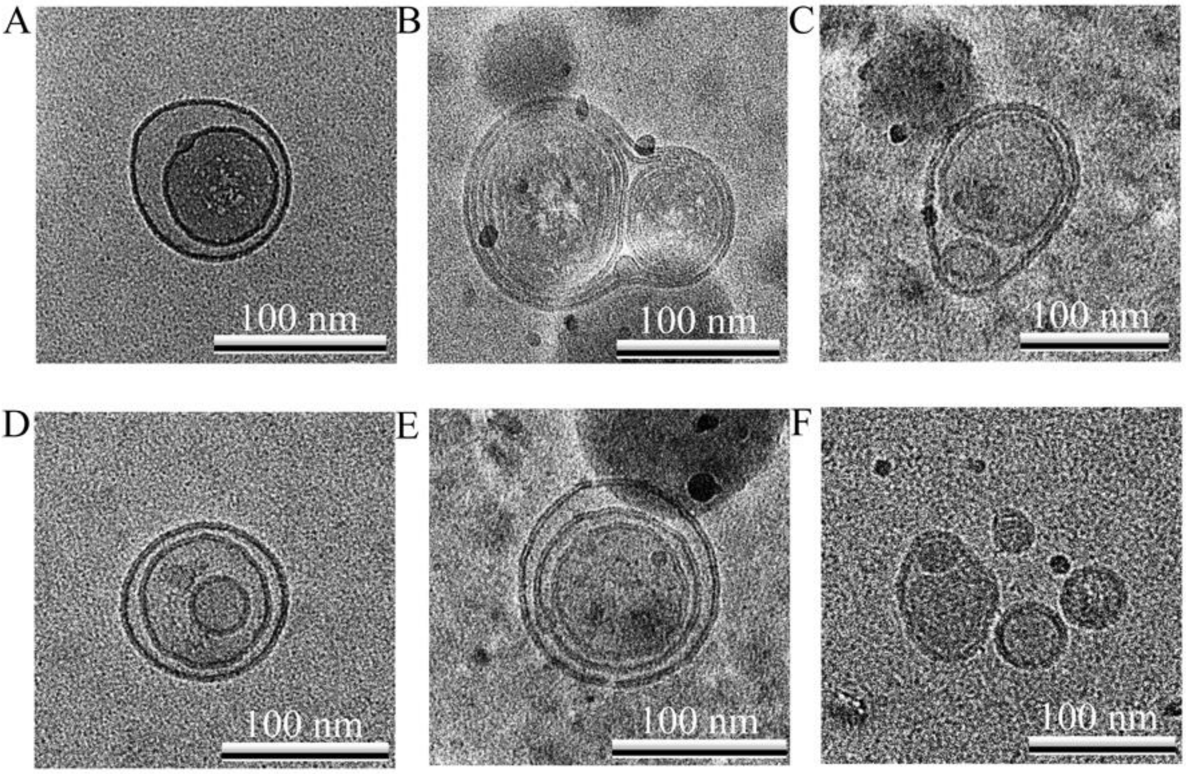


**Figure S6: Particles formed after prolonged ethanol treatment.** On the fourth day following ethanol treatment, the sample was filtered through 0.22 micron syringe filter and subjected to ultracentrifugation (at the same speed used to precipitate yeast ribosome) and viewed in cryo-TEM. Particles of different sizes (<200nm in size) with unusual structural features were seen (A-F).

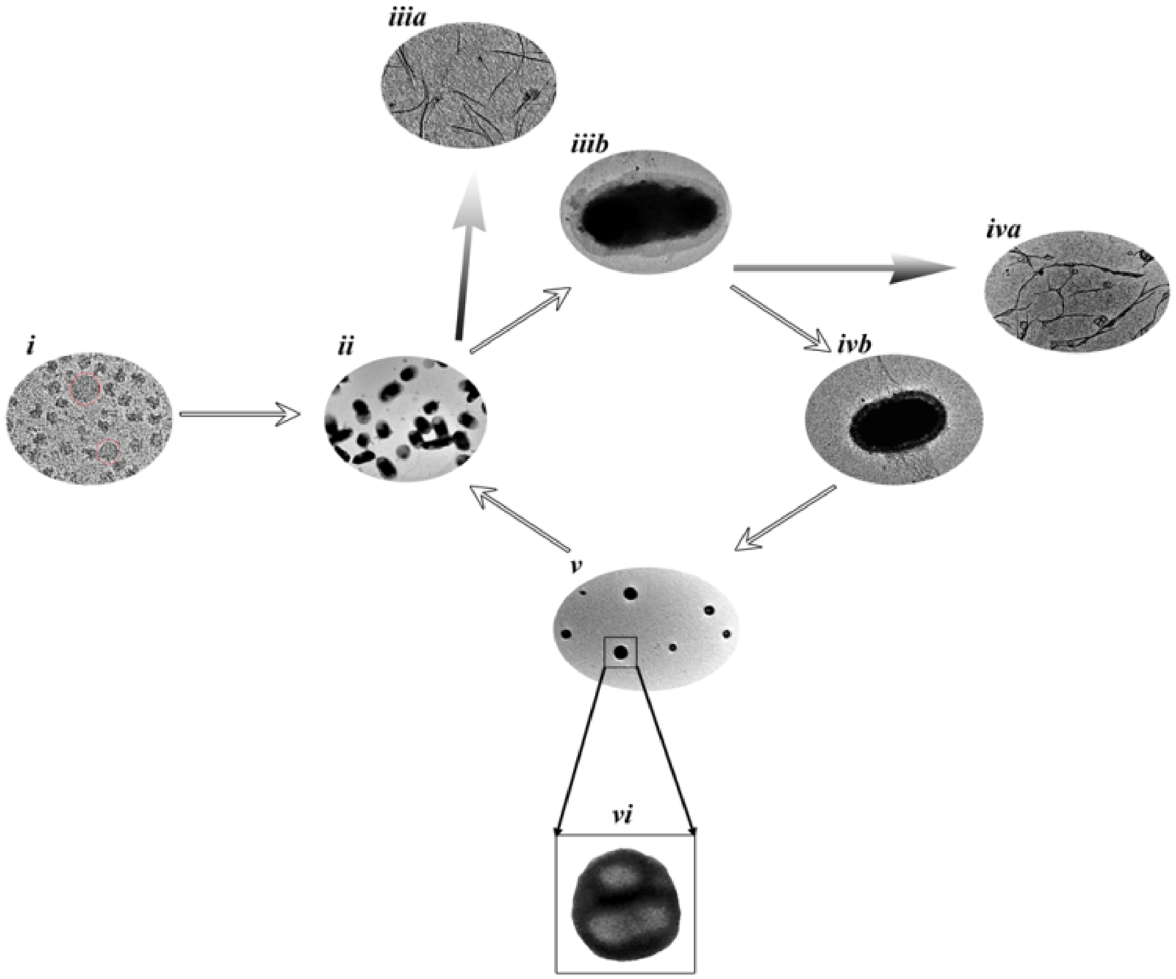


**Figure S7: Schematic representation of the life cycle of maturation of the bacterial species.** Initially spherical structures were seen in the Yeast ribosome preparation (i). Formation of larger spores/vegetative cells occurred when Yeast ribosomes were added directly to culture media (ii). Upon overnight salt-ethanol stress followed by centrifugation supernatant revealed fibrils (iiia) while shrunken cells were found in pellet (iiib). Continued salt-ethanol stress produced more fibrils in the supernatant (iva) and smaller cells in pellet (ivb). Nano-spores were eventually visible when the supernatant was filtered through 0.22 micron syringe filter following prolonged salt-ethanol stress treatment (v). Close up view of TEM image of a nano spore (vi). The nano-spores (formed under salt-ethanol stress), when cultured in LB, again transformed to bacterial cells (ii).
